# Deciphering Global Patterns of Marine Microbial Community Assembly and Network Stability

**DOI:** 10.1101/2025.09.19.677047

**Authors:** Pranathi Ravikumar, Aarti Ravindran, Karthik Raman

**Affiliations:** Department of Biotechnology, Bhupat and Jyoti Mehta School of Biosciences, Indian Institute of Technology Madras, Chennai, 600036, Tamil Nadu, India; Centre for Integrative Biology and Systems mEdicine (IBSE), Wadhwani School of Data Science and AI, IIT Madras, Chennai, 600036, Tamil Nadu, India; Department of Data Science and AI, Wadhwani School of Data Science and AI, Indian Institute of Technology Madras, Chennai, 600036, Tamil Nadu, India

**Keywords:** marine microbiome, community assembly, co-occurrence networks, microbial diversity, network stability

## Abstract

Microbial community assembly processes can help us understand ecology, evolution, and climatic influences on community composition while presenting opportunities for biotechnological applications and improved marine conservation. In this study, we have investigated species richness patterns, community assembly mechanisms, and interaction patterns of marine microbial communities by analysing 16S rRNA amplicon sequencing data from 4,611 samples collected from major ocean microbiome projects. Using neutral community models, the iCAMP framework, and co-occurrence network analyses, we showed that stochastic processes drive microbial community assembly across all latitude zones. Drift was the primary driver of community assembly in polar communities. The polar microbial communities exhibited the highest modularity and network robustness, but were more vulnerable to hub removal. While previous studies have attributed higher stability of the polar communities to environmental filtering, our analyses reveal that the resilience of the community is dependent on a few central taxa. By classifying the genera as generalists and specialists, we further highlight the role played by the specialist taxa in maintaining the stability of the marine microbial community, especially under the pressures of climate change and global warming. Overall, our findings offer a latitudinal perspective on ocean microbial community stability and the effect of anthropogenic disturbances on these communities.

## 1 Introduction

Microbial community assembly processes explain the fundamental principles of ecology, evolution, and climatic variations and can be used for biotechnological, environmental applications, and management. Large-scale ocean microbiome projects, including the Tara Oceans Project [1], Malaspina Expedition [2, 3], Global Ocean Sampling [4, 5], and Bio-GO-SHIP [6], have revolutionised our understanding of microbial life in the oceans. These initiatives have enabled public access to microbial community data across spatial and environmental gradients, supporting global-scale analysis of the ocean microbiome.

To investigate the ecological mechanisms underlying community assembly, researchers employ conceptual frameworks and statistical methods that differentiate between deterministic (niche-driven) and stochastic (neutral) processes [7–9].Deterministic processes such as environmental filtering and species interaction structure communities by selecting traits suited to particular environmental conditions. On the other hand, stochastic processes include ecological drift, dispersal limitation, and historical contingencies resulting in community patterns that arise from random events and colonisation. Models such as the Neutral Community Model [10] and the iCAMP (integrated Community Assembly Mechanisms Profiling) framework [11] enable quantitative estimation of the contribution of these processes in driving community assembly.

While community assembly models uncover abiotic drivers of community composition, they cannot capture the interactions of taxa within the communities. Hence, co-occurrence network analysis can be a powerful tool in microbial ecology. Co-occurrence networks are constructed by identifying statistically significant correlations between taxa across samples, enabling the inference of potential ecological interactions [12]. The structure of these networks can be quantified using network properties such as modularity, degree distribution, clustering coefficient, and natural connectivity. These properties can reveal emergent properties such as stability, robustness, and keystone taxa in the communities [13].

These network topologies often explain the ecological roles played by the taxa. For example, some genera may act as generalists and have high prevalence across environments. On the other hand, they may be specialists, having niche-specific prevalence. These patterns can be used to understand how microbial ecosystems respond to perturbations such as warming, acidification, and nutrient fluctuations [14, 15].

One of the most widely used methods to characterise marine microbial communities in such studies is 16S rRNA gene amplicon sequencing [16–18]. This technique targets conserved regions of the ribosomal rRNA gene to profile taxonomic composition at relatively low costs and high throughput [19]. It can be used to estimate the microbial diversity and biogeographic patterns to understand ecological processes in these communities [20].

We have utilised 16S rRNA amplicon sequencing data from ocean microbiome projects in this study. We have investigated the ecological mechanisms governing community assembly processes in the ocean microbial community using the Sloan Neutral Community Model (NCM) and the iCAMP framework. Furthermore, co-occurrence networks were constructed and have been used to uncover structural features and keystone taxa. The microbes in these networks were classified as generalists and specialists based on their prevalence to evaluate their topological roles and contributions to network robustness.

Through this integrative framework, the study aims to provide a comprehensive understanding of the ecological processes structuring global marine microbial communities, focusing on how latitude-mediated environmental gradients influence the presence of taxa within these communities, and how they interact and function as a collective ecological network.

## 2 Methods

### 2.1 Data Retrieval And Preprocessing

A total of 4611 16S amplicon sequencing samples were obtained from multiple ocean microbiome projects, including Tara Oceans Project [1], Malaspina Expedition [21], Global Ocean Sampling [4] and Bio-GO-SHIP [6]. Additionally, marine samples from the Australian Microbiome Project and the Earth Microbiome Project were incorporated [22, 23].The number of samples taken from each of these projects has been mentioned in Supplementary Table 1. The sample locations have been inferred from the latitude and longitude data and incorporated into the metadata file (Supplementary File 1 and depicted in Supplementary Figure 1.

The quality filtering and preprocessing of sequence data were performed using the Quantitative Insights into Microbial Ecology (QIIME2) pipeline (version 2024.5) [24]. The sequences were clustered using the DADA2 algorithm [25]. The representative sequences from each ASV were aligned to the GreenGenes2 reference database (v22.10) [26], and taxonomic classification was performed using the RDP Classifier. The unclassified ASVs have been filtered out from further downstream analysis. The parameters used in the denoising process and the primers used for training the RDP Classifier are mentioned in Supplementary File 2. The results of the denoising process are described in detail in Supplementary File 3.

### 2.2 Statistical Analysis

The Chao1 and Shannon Diversity indices were calculated using the ‘vegan’ package v2.6.10 [27] in R Statistical Computing Environment [28]. The Kruskal-Wallis test was performed using the ‘Phyloseq’ v.1.52.0 package [29] to test the significance of the differences in Shannon Diversity metrics amongst samples. Using the ‘adonis2’ function available in the ‘vegan’ package, Permutational Multivariate Analysis of Variance (PERMANOVA) was performed to identify the significant factors contributing to the variation between samples. The Bray-Curtis dissimilarity index was used as the basis for the Principal Coordinate Analysis (PCoA), and was then used to plot the beta diversity of the samples. Beta dispersion, which shows the spread or variability in the community composition, was estimated using the ‘betadisper’ function. The Canonical Correspondence Analysis (CCA) and Variation Partition Analysis (VPA) were performed using the ‘vegan’ package to corroborate the PERMANOVA results.

### 2.3 Community Assembly

The Neutral Community Model (NCM) was used to estimate the contribution of stochastic and deterministic processes in community assembly of marine samples [10]. The ‘MicEco’ and ‘vegan’ packages were used for the analysis at the genus level without any filtering criteria applied. Another approach used to reveal the patterns of stochasticity and determinism and their influence on microbial communities was to calculate the Levin’s Niche breadth (B) index (Bcom) [30]. This analysis was conducted using the ‘niche.width’ function in the “spaa” package available in R [31]. NST was calculated using the NST package in R (v.3.1.10) [32]. This was carried out with a cutoff of at least 50 reads in a sample and a taxa prevalence of 0.1 per cent. A phylogenetic-bin-based framework (iCAMP) was used to infer community assembly mechanisms. A cutoff of at least 50 reads and a minimum prevalence of 0.1% was used to make it computationally less intensive without losing too much information. The icamp.big, icamp.bins and icamp.boot functions in the R package iCAMP function were used [11].

### 2.4 Co-occurrence Networks Construction And Analyses

The microbial co-occurrence network was constructed based on the association values computed using SPIEC-EASI (SParse InversE Covariance Estimation for Ecological Association Inference) [12]. A cutoff of at least 50 reads and a minimum of 1% prevalence was used to filter out noise and avoid spurious interactions. Network analysis was performed using the ‘igraph’ [33] and ‘NetCoMi’ package [34]. The plots generated in the study have been generated using the ‘ggplot’ package [35].

## 3 Results

### 3.1 Multiple Factors Contribute To Variation In The Marine Bacterial Community

The metadata uniformly available across all datasets included latitude and longitude coordinates, the ocean (determined from these coordinates), and the sample origin, categorised as marine, coral-associated, sponge-associated, or marine sediment. To capture the differences in the community composition, Bray-Curtis dissimilarity was calculated. PERMANOVA was performed to quantify how much these factors contribute to the variation in diversity. PERMANOVA results showed that the latitude zones explain 12.5% and the oceans (which are correlated with the latitude) explain 11.9% of the variance (Supplementary Table 2). The Bray-Curtis distance was ordinated with PCoA. The first two principal axes accounted for more than 30 per cent of the variation (Supplementary Figure 2). The bacterial communities were clustered into three groups based on the latitude zones (Polar, Temperate and Tropical, Figure 1A). The beta dispersion of samples was evaluated to compare microbiome composition variability across different latitude zones. Samples from temperate regions exhibited the highest beta dispersion, followed by those from tropical and polar regions (Figure 1B). The differences in beta dispersion between latitude zones were statistically significant (Kruskal-Wallis test *p-value<* 0.001).

**Fig. 1:**
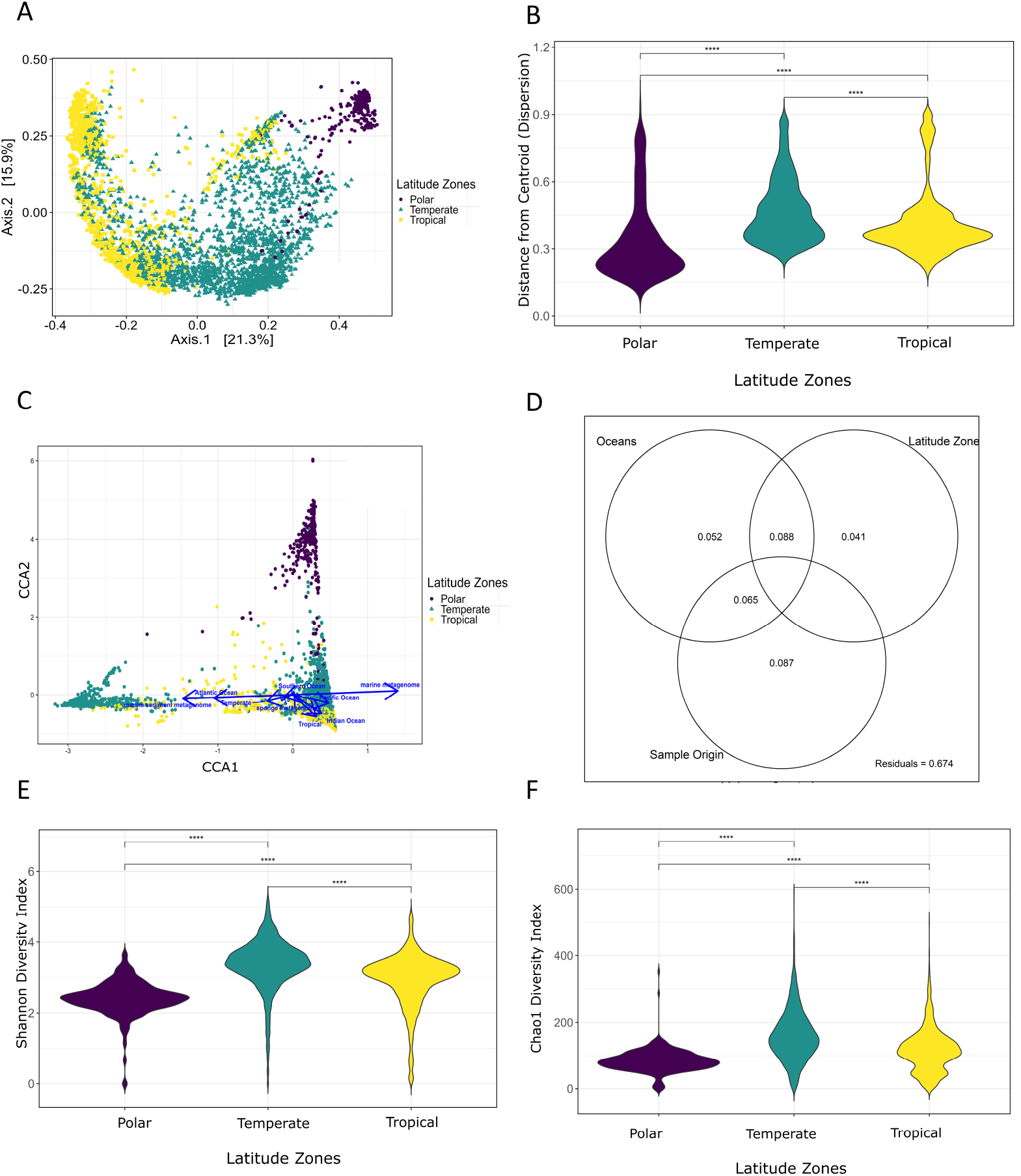
Compositional Analysis of Marine Microbial Communities. A: PCoA. B: Beta-dispersion (Kruskal– Wallis test, *p-value<* 0.001). C: CCA depicting the effect of different environmental factors on the Bray–Curtis distance. D: VPA. E: Shannon diversity of samples grouped by latitude zones. F: Chao1 diversity of samples grouped by latitude zones.

CCA revealed that sample origin, latitude zones, and ocean significantly influenced marine microbial community composition. (Figure 1C). The analysis showed that latitude zones, sample origin and the ocean of origin had significant correlations (*p-value<* 0.05) with the bacterial communities (Supplementary Table 3). VPA was performed to determine how much variation in species composition can be uniquely and jointly explained by different environmental variables. The ocean of origin was the most significant factor influencing marine microbial community composition among the tested variables, followed by latitude (Figure 1D).

Alpha diversity indices (Shannon and Chao1) were also calculated to complement the beta diversity patterns. ANOVA results and effect size analysis indicated that latitude zones are the primary contributors to differences in alpha diversity indices (Supplementary Table 4). The sample origin and the ocean of origin also showed significant differences in all alpha diversity indices. Polar regions showed the lowest Shannon and Chao1 diversity indices (Figure 1E, F). The bacterial communities present in the temperate regions had the highest alpha diversity indices (Shannon diversity index: 3.37 ± 0.72 and Chao1 diversity index: 160.01 ± 76.71). Pairwise Wilcoxon tests were performed, which showed significant differences in the alpha diversity indices between polar, temperate and tropical zones (Figure 1E, F). Given the high correlation between ocean and latitude, and to avoid collinearity in downstream analyses, we retained latitude as the primary environmental variable

The Kruskal-Wallis test was conducted at the phylum level to assess taxonomic shifts across latitude zones. *Campylobacterota, Bacteroidota* and *Firmicutes* showed the most pronounced differences in abundance (Supplementary Table 5). *Campylobacterota* was more abundant in temperate regions, whereas *Bacteroidota* and *Firmicutes* were enriched in polar and tropical regions, respectively. Genus-level agglomeration retained an optimal number of ASVs while providing adequate resolution for our analysis. *Pelagibacter* was the most abundant genus in the ocean microbial community.

### 3.2 Marine Communities Display High Stochastic Community Assembly Processes

The NCM analysis was employed to assess the role of neutral processes, such as random dispersal and ecological drift, in shaping the assembly of microbial communities (Figure 2A-C). The microbial communities in the polar regions had the lowest R2, and the temperate regions had the highest R2, indicating the best fit with the neutral model. The estimated dispersal parameter (Nm) reflects the combined influence of migration and community size on taxon distribution and was highest in the polar bacterial communities.

**Fig. 2:**
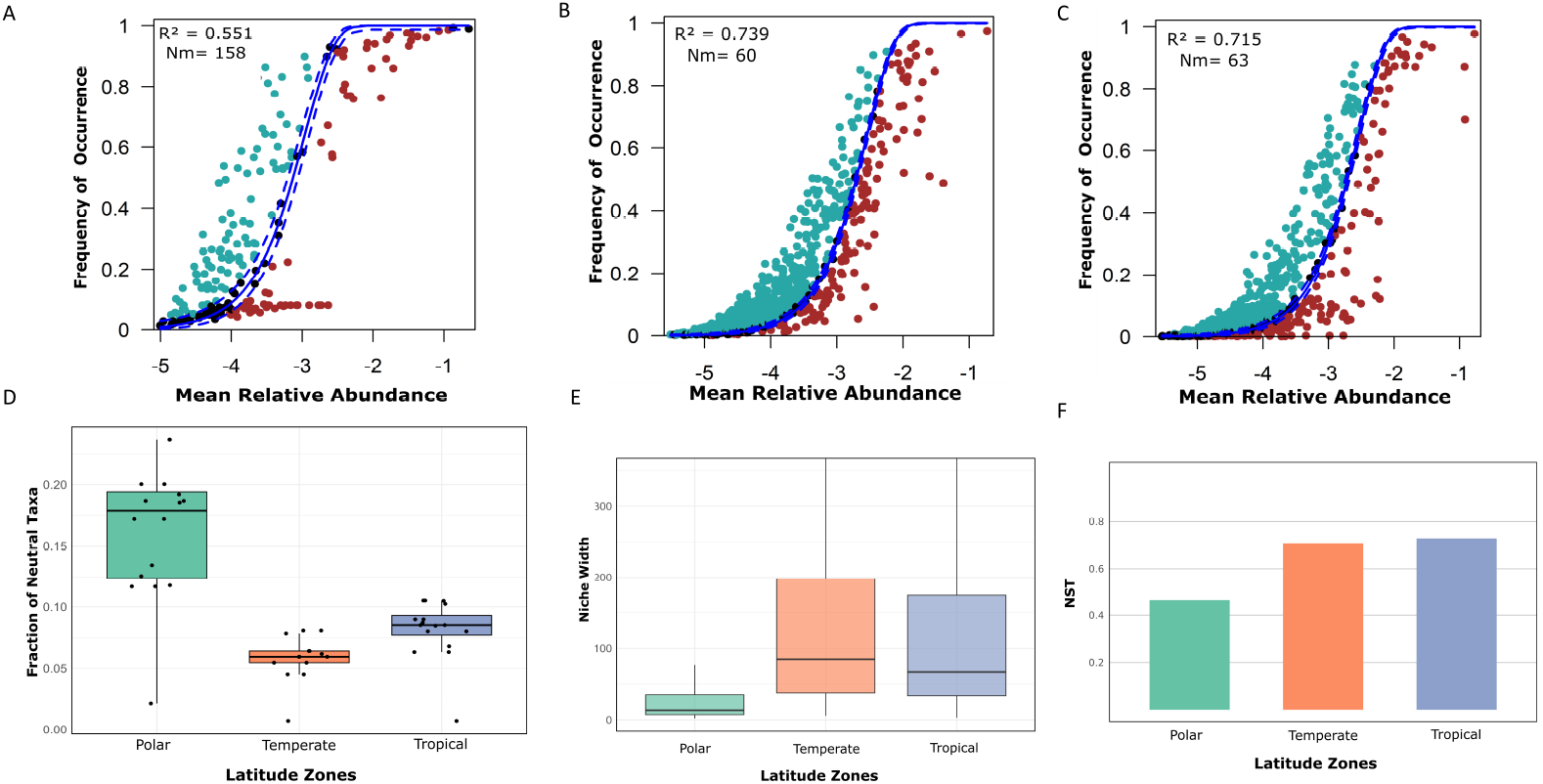
Assessment of neutral model fit across marine microbial communities in polar, temperate, and tropical zones. (A–C) Comparison of the fit of polar (A), temperate (B), and tropical (C) marine microbial communities to the neutral community model. The blue line represents the predicted distribution under the neutral model. Taxa colored blue fit the model predictions, while green and red indicate taxa with significantly higher or lower abundances than expected, respectively. (D) Fraction of neutral taxa identified in polar, temperate, and tropical marine regions. (E) BCom values illustrating the niche width of taxa across polar, temperate, and tropical communities. (F) NST values of the polar, tropical, and temperate regions.

Following the NCM analysis, the fraction of taxa that conformed to neutrality was quantified to assess if stochastic processes could explain the community composition. There was a significant difference in the fraction of taxa conforming to NCM across latitude zones (Figure 2D) (Wilcoxon test: *p-value <* 0.001). The polar and temperate regions exhibited the highest and lowest fraction of neutral taxa, respectively.

A null model analysis was conducted on PERMDISP to examine the variation in community composition across latitude zones. The null model test results showed that dissimilarities in the polar regions were significantly lower than null expectations (Supplementary Table 7). The temperate regions had a higher dissimilarity than the null expectations, suggesting that environmental filtering plays a strong role in polar microbial community formation and a weaker role in the temperate regions.

We also compared the niche width across latitude zones to assess the factors driving community turnover. Polar communities displayed lower BCom values (Figure 2E), indicating that the community composition is relatively similar across the samples. In contrast, temperate and tropical regions displayed significantly broader BCom, suggesting greater beta-diversity and spatial turnover. NST was calculated to quantify the relative contribution of the stochastic and deterministic processes (Figure 2F). These results imply that environmental heterogeneity and niche-based processes influence the community composition in tropical and temperate regions, whereas the polar communities are more homogeneous.

We further applied the iCAMP framework to partition stochastic and deterministic processes into specific ecological mechanisms such as selection, dispersal, and drift to understand the ecological processes driving community assembly (Figure 3A-C). Drift was the dominant driver of community assembly across all latitude zones, with its influence being particularly pronounced in the polar regions. Deterministic processes, such as homogeneous and heterogeneous selection, played a relatively more important role in the tropical and temperate communities than in the polar communities.

**Fig. 3:**
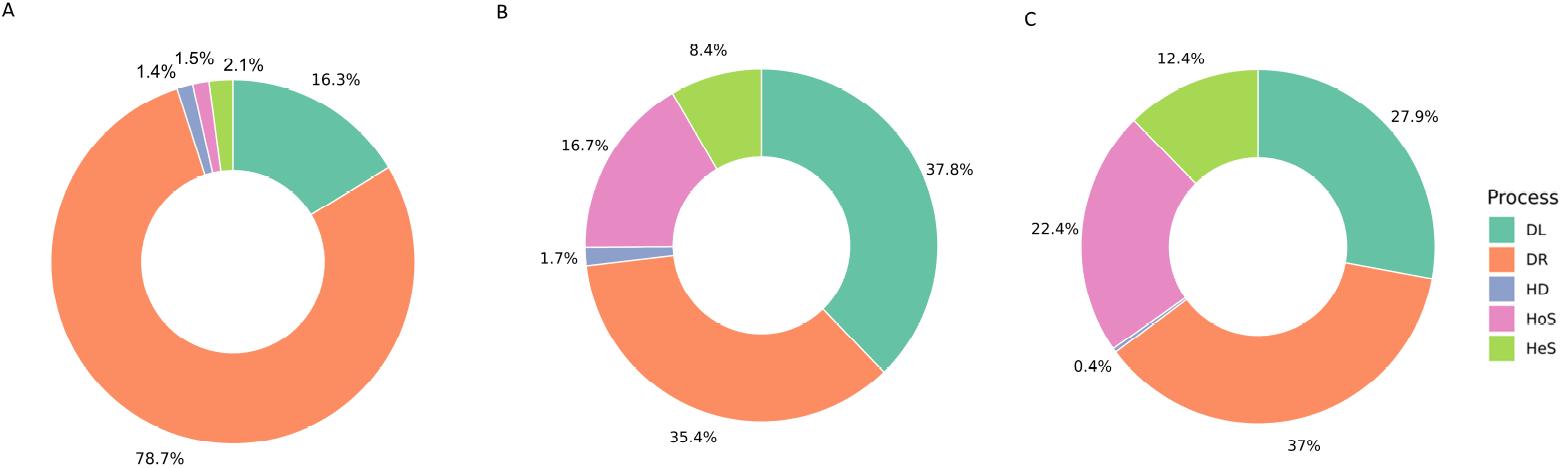
Patterns of neutrality, niche breadth, and community assembly processes across latitudinal marine microbial communities. (A–C) Relative contributions of community assembly processes—including dispersal limitation (DL), drift (DR), homogeneous selection (HoS), and heterogeneous selection (HeS)—in (A) polar, (B) temperate, and (C) tropical regions.

These results reveal that microbial community assembly in the ocean is shaped by stochastic and deterministic processes, with strong environmental filtering playing dominant but latitude-dependent roles.

#### 3.3 Network Modularity Enhances Perturbation Resistance In Polar Microbiomes

We then proceeded to study the co-occurrence patterns using network analysis to understand the ecological interactions shaping these communities. The network constructed using the above methods had 554 nodes and 4227 edges. Overall, positive interactions predominate (92.81%) in the co-occurrence networks. After removing the negative edges, the modularity of the network was calculated using the Louvain algorithm [36]. The network has a modularity of 0.66. Given the modularity value (*>* 0.5), the network has an overall modular structure [37] with 10 sub-modules with sizes varying from 26 to 101 nodes (Supplementary Figure 3). The network has an average degree of 15.26 and an average clustering coefficient of 0.218 (Supplementary Figure 3).

Nodes in the co-occurrence network largely belonged to the phyla *Proteobacteria* (44.7%), *Bacteroidota* (17.15%), *Actinobacteriota* (3.79%), *Verrucomicrobiota* (3.61%) and *Chloroflexota* (3.07%) (Figure 4A). Inter-phyla interactions comprised 2569 of the total interactions, with the remaining 1354 occurring within phyla. Inter-phyla interactions vastly outnumber intra-phyla interactions across all phyla. *Proteobacteria* and *Thermoplasmota* exhibited the highest inter-phyla to intra-phyla interaction ratios (Figure 4B). *Proteobacteria* and *Bacteroidota* accounted for 72.2% and 15.01% of all the intra-phyla interactions (Figure 4C) and had the largest number of inter-phyla interactions (Figure 4D).

**Fig. 4:**
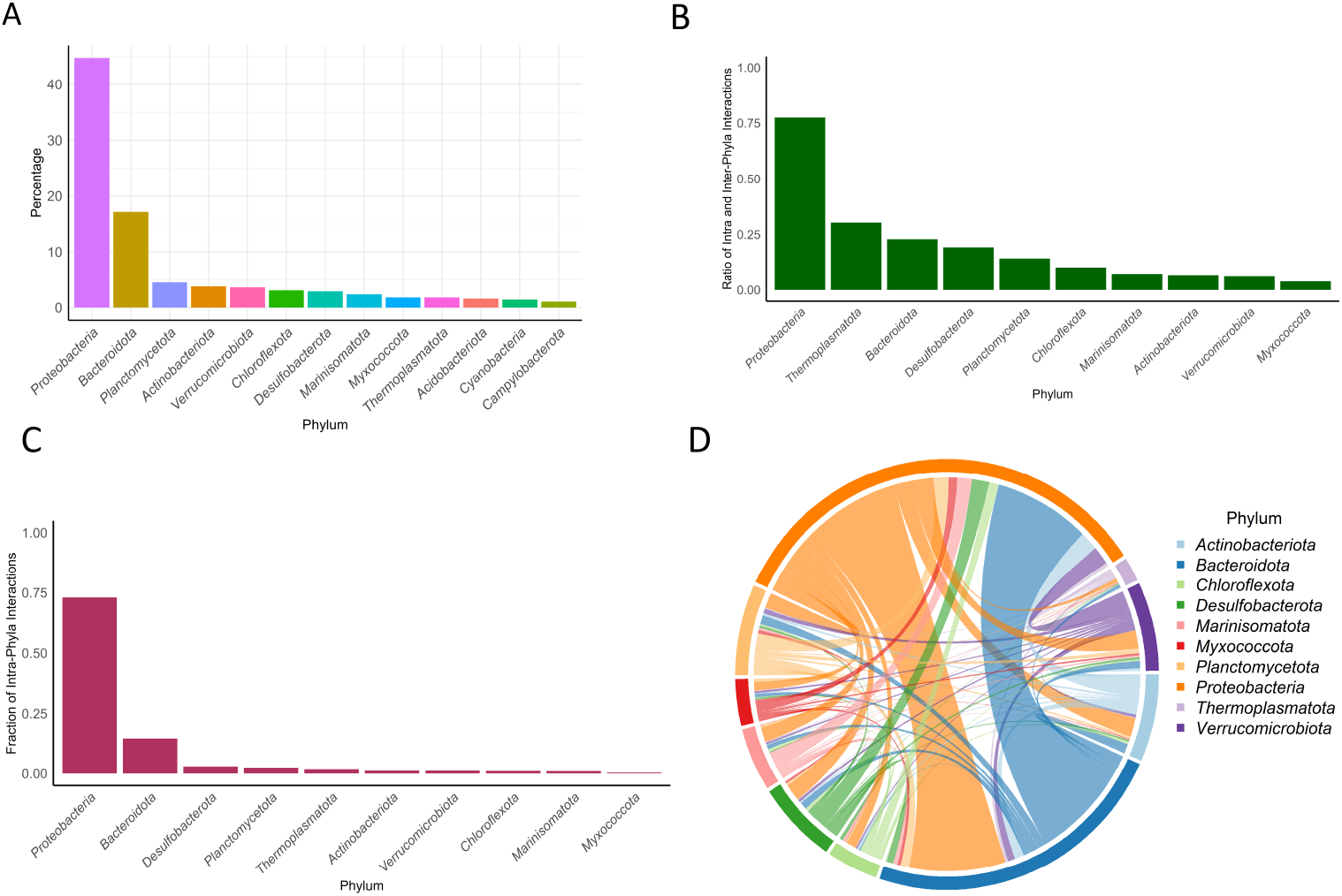
Phylum-level structure and interaction patterns in the ocean microbial community network. (A) Distribution of phyla represented by nodes within the ocean microbial community network. (B) Ratio of inter-phyla to intra-phyla interactions among nodes, illustrating connectivity between and within different phyla. (C) Proportion of intra-phyla interactions attributed to each phylum, highlighting internal network structure. (D) Composition and types of inter-phyla interactions across the entire ocean microbial network.

We then constructed latitude-zone-specific co-occurrence networks (Figure 5A-C). The polar, temperate and tropical co-occurrence networks had 201, 630 and 365 nodes (Table 1). The degree distribution of these networks is shown (Supplementary Figure 3A-C). The temperate co-occurrence network had the highest average degree of 15.81. The co-occurrence network constructed using the samples from polar regions had the least variability in the degree (Figure 5D). To assess network robustness, we removed high-degree nodes (hubs) and evaluated the resulting changes in natural connectivity, a measure of network resilience. Natural connectivity decreases with increasing percentage of nodes removed (targeted by degree) across all three networks. The slopes of this decline were 0.00060 for polar networks, 0.00042 for temperate networks, and 0.00055 for tropical networks. A less negative slope in the temperate network indicates greater robustness to targeted node removal than the polar and tropical networks. The polar network had the lowest natural connectivity and the steepest drop upon removing nodes of the highest degree (Supplementary Figure 4). The polar co-occurrence network had the highest density (Table 1). The clustering coefficient was highest in the tropical co-occurrence network. The polar, temperate and tropical co-occurrence networks across latitude zones have a modular structure, with 14, 11 and 12 sub-modules, respectively [37]. The modularity is reported to be the highest for the polar co-occurrence network.

**Table 1:** Properties of polar, temperate and tropical microbial co-occurrence networks, using taxa which are prevalent in at least one per cent of the samples

| Property | Polar | Temperate | Tropical |
| --- | --- | --- | --- |
| Number of Nodes | 198 | 630 | 365 |
| Number of Edges | 483 | 4908 | 1798 |
| Network Density | 0.025 | 0.025 | 0.027 |
| Average Degree | 6.328 | 15.581 | 9.852 |
| Clustering Coefficient | 0.225 | 0.215 | 0.227 |
| Modularity | 0.778 | 0.691 | 0.763 |
| Ratio of negative to positive edges | 0.123 | 0.056 | 0.183 |

**Fig. 5:**
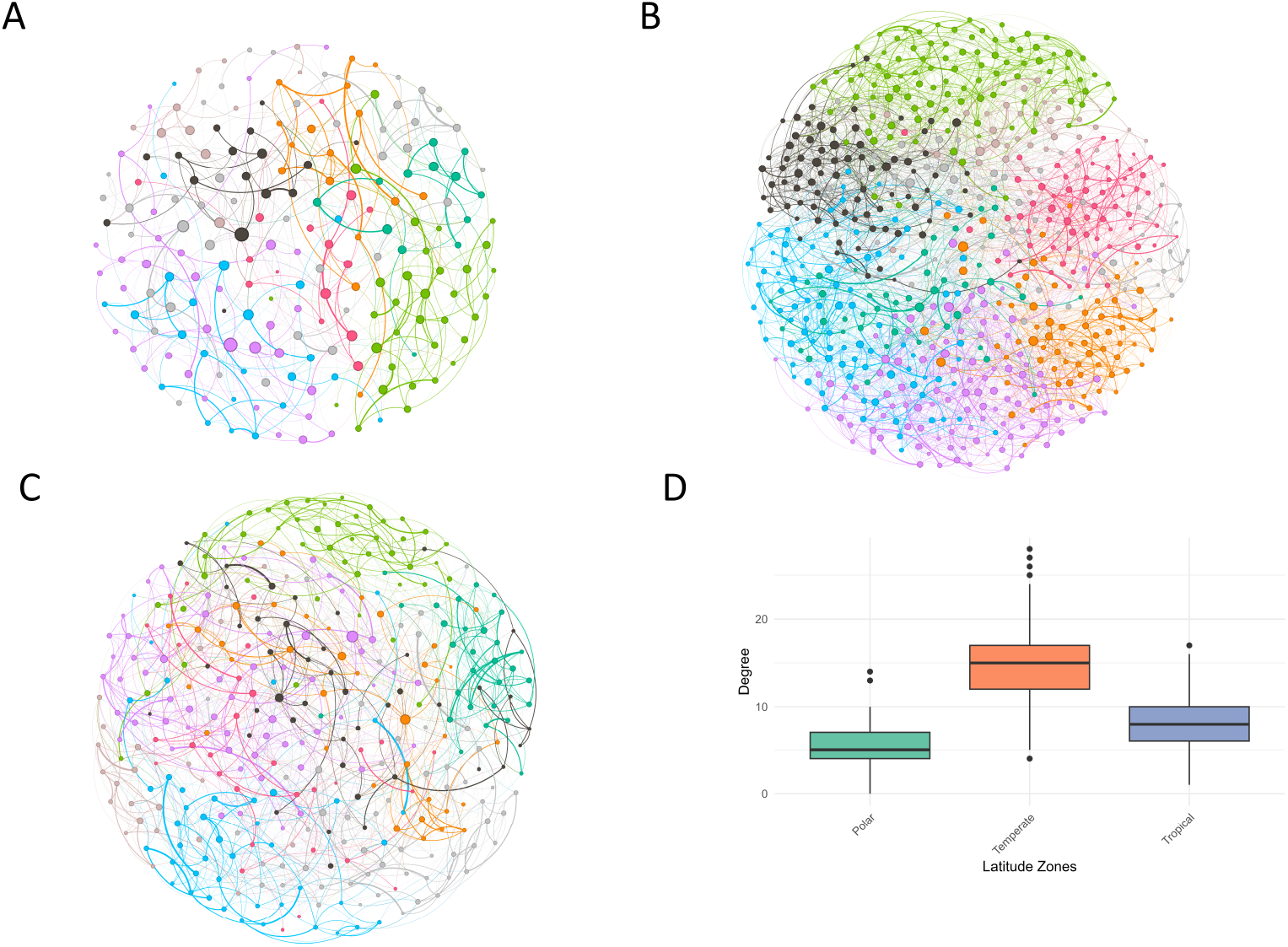
Modular structure and connectivity of microbial co-occurrence networks across polar, temperate, and tropical marine regions. (A–C) Co-occurrence networks of polar (A), temperate (B), and tropical (C) marine microbial communities, with nodes colored according to their respective network modules, highlighting modular organisation within each region. (D) Comparison of the average degree (network connectivity) across polar, temperate, and tropical microbial co-occurrence networks, illustrating variability in community connectivity among regions.

In addition to the global network properties, the topological properties of the individual genera help understand their ecological roles. Genera are classified into four buckets (Figure 6) based on their within-module (Z) and among-module connectivity (P) [38]. The peripherals (low Z and low P) are genera that have fewer interactions with other genera within their modules. Most genera in the network were peripherals. The module hubs (high Z and low P) contain genera that have a higher number of interactions with other genera within their own modules. The polar, temperate, and tropical co-occurrence networks had 3, 4, and 1 module hubs, respectively. The polar co-occurrence network had the largest percentage of connectors (high P, low Z) compared to the other co-occurrence networks. Genera acting as connectors in the polar microbial community include *Moritella, Oleispira*, and *Marinomonas*.

**Fig. 6:**
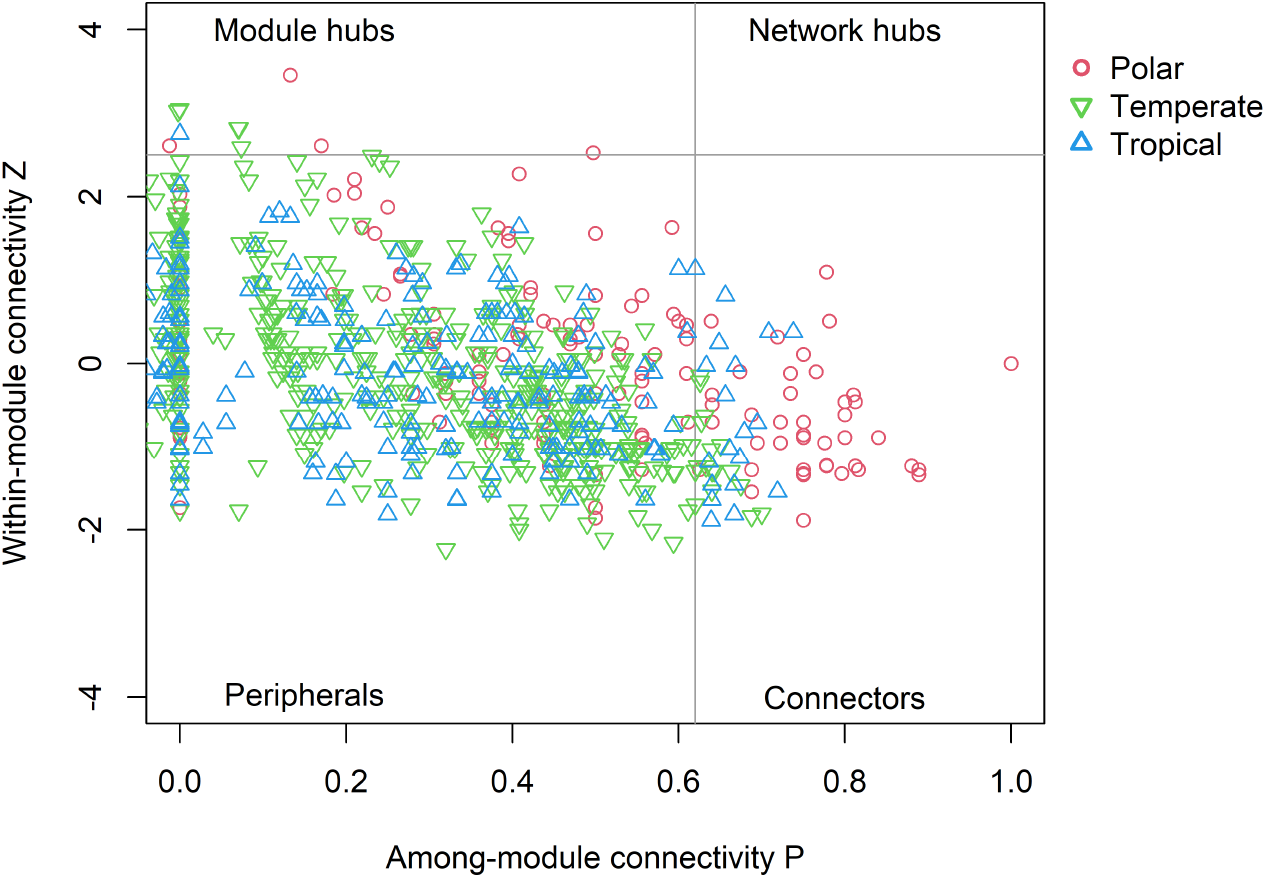
Role of taxa in polar, temperate, and tropical marine microbial communities. Taxa are classified according to their network roles—peripherals, connectors, module hubs, and network hubs— based on patterns of inter- and intra-module connectivity within each co-occurrence network. This highlights the structural and functional positions of taxa across distinct climatic regions.

### 3.4 Specialist Taxa Play A Major Role In Maintaining The Overall Structure Of The Ocean Microbiome

Genera are classified as marine generalists (prevalence *>* 50%) and specialists (prevalence *<* 5%) (Figure 7A). Under this criterion, 8% of the genera were classified as generalists and 12% of the genera were classified as specialists. Generalists and specialists tend to interact amongst themselves, rather than with other genera (Figure 7B). Specialists show a larger variability in the degree compared to the generalists (Figure 7C). The average degrees of the generalists and specialists were 14 and 13 with standard deviations of 2.85 and 4.11, respectively.

**Fig. 7:**
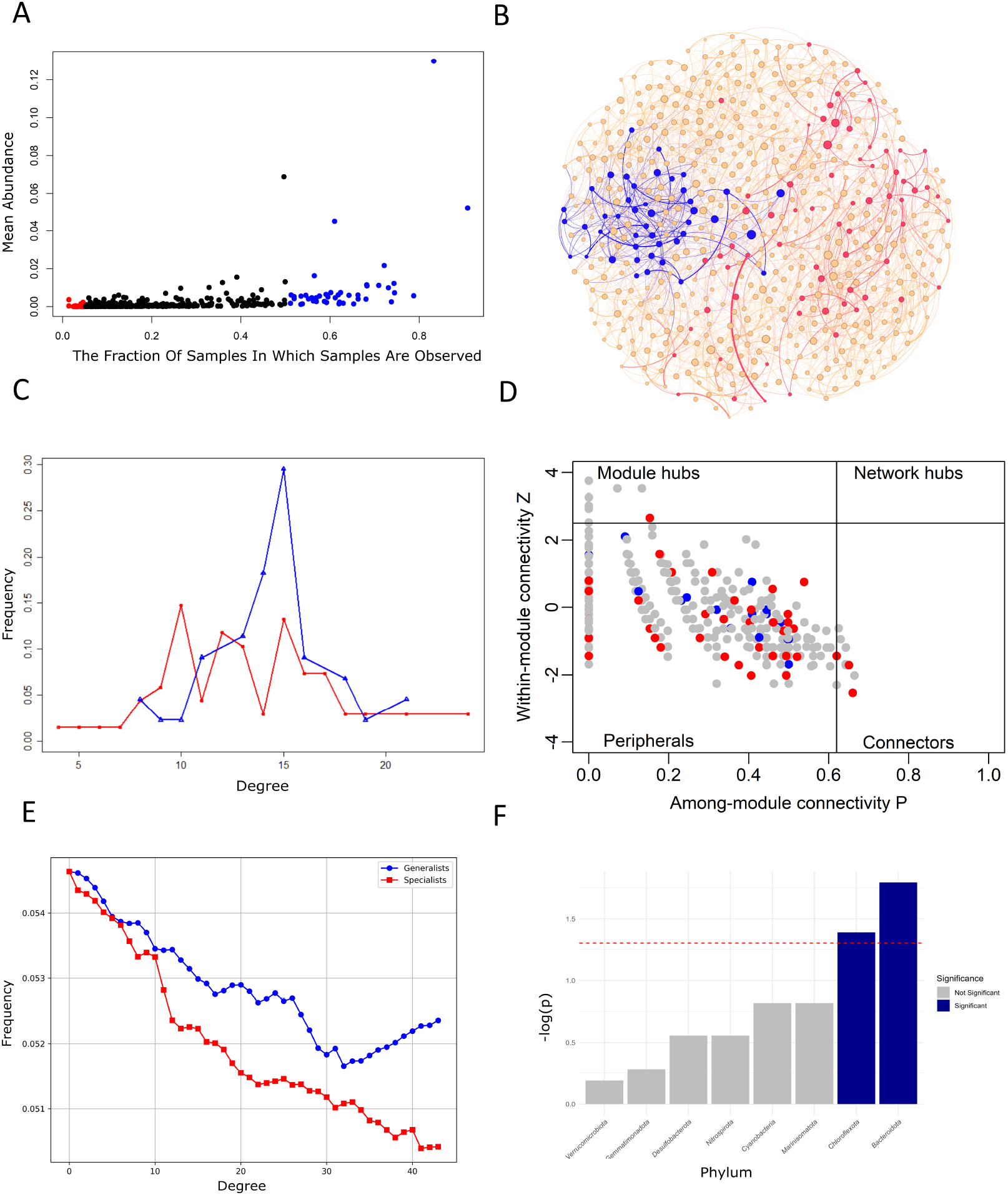
Analysis of the roles of the generalists and specialists in ocean microbial communities. (A) Occupancy plot depicting the mean abundance and prevalence of different taxa in the ocean microbial network. (B) Overall ocean microbial co-occurrence network depicting the interactions between generalists and specialists. (C) Degree distribution of the generalist and specialist taxa in the ocean microbial network. (D) Topological roles of the generalist and specialist taxa in the ocean microbial network based on their inter-module and intra-module connectivity. (E) Changes in the natural connectivity of the overall ocean microbial network after removing specialists and generalists to highlight their importance. (F) Phyla with a significantly different number of generalists and specialists in the ocean microbial co-occurrence network.

To further understand the ecological roles of the generalists and specialists, the inter-module and intra-module connectivity were calculated and plotted (Figure 7D). Specialists act as module hubs and connectors. On the other hand, generalists have a limited role in the co-occurrence network as peripherals. To determine the role of the generalists and specialists on network stability, the natural connectivity of the network was calculated after progressively removing the generalist and specialist nodes of the highest degree. There is a steep drop in the natural connectivity of the network upon removal of the specialists compared to removal of the generalists (Figure 7E).The generalists belonged to phyla *Proteobacteria, Bacteroidota, Actinobacteriota, Cyanobacteria, Mariniso-matota* and *Verrucomicrobiota*. The specialists belonged to a broader range of phyla, including *Proteobacteria,Chloroflexota, Bacteroidota, Actinobacteriota, Desulfobacterota, Nitrospirota*, and *Verrucomicrobiota*. There was a significant difference in the number of generalists and specialists belonging to the phyla *Chloroflexota* and *Bacteroidota* (Figure 7F) (Fisher test *p-value <* 0.05). Approximately 67% of the generalists belonged to *Bacteroidota*; in comparison, only 26% of the specialists belonged to the same phylum. On the other hand, 30% of the specialists belonged to *Chloroflexota* and no generalists were identified in this phylum. Specialists had significantly higher inter-phyla interactions than intra-phyla interactions (Fisher test *p-value <* 0.005). Generalists have nearly equal inter- and intra-phyla interactions (Supplementary Figure 5).

## 4 Discussion

We structured our study on the latitude variation as it provided the most homogeneous distribution of samples while capturing climatic and environmental gradients. Previous studies have also identified that latitudinal gradients strongly affect the diversity in marine bacterial communities [39–41], which is consistent with the intermediate disturbance hypothesis, which proposes that diversity peaks when environmental disturbances occur at moderate levels [42].VPA suggests that the sample origin is a major contributor to bacterial composition. Given that only a small fraction of samples (171 out of 4,583) in our dataset are host-associated, this factor was not detected as significant in the PERMANOVA analysis. Overall, the three factors, namely latitude, sample origin, and ocean, are highly interrelated variables, and the discrepancies in the batch sizes render meaningful insights challenging.

Diversity analysis at the genus and phylum levels was performed to understand the community composition. *Pelagibacter* is the most dominant genus in the marine samples, consistent with its known role as a key player in marine microbial ecosystems [43]. The differential abundance of *Bacteroidota, Firmicutes* and *Campylobacterota* across latitude zones implies that specific phyla may adapt to specific environmental conditions (Supplementary Table 5). *Bacteroidota* is enriched in polar regions because they can efficiently utilise complex carbohydrates using fermentation pathways in energy-limited environments, providing a competitive advantage in lower temperatures [44]. *Firmicutes* have the highest abundance in tropical regions through endospore formation, producing highly resistant spores that can withstand heat and UV radiation [45, 46]. *Campylobacterota* thrive in temperate regions because their key autotrophic enzymes, such as ATP citrate lyase, exhibit mesophilic temperature optima, aligning with the moderate thermal regimes of temperate marine waters [47]. While this approach sheds light on key ecological mechanisms, the utilisation of 16S rRNA data, results in a lack of species-level resolution.

The relative contributions of the deterministic and stochastic processes were estimated across different latitude zones. The host-associated samples and the marine sediment samples have been removed from this analysis to reduce the confounding factors. The neutral theory model, null theory model and iCAMP frameworks were used to understand the roles of selection, dispersal and environmental filtering in shaping the structure of the marine microbial communities [11, 30]. These three approaches integrate taxonomic community composition, dispersion metrics and phylogenetic analyses, respectively, to provide a holistic understanding of marine microbial community assembly across latitude zones.

NCM analysis revealed that the dispersal parameter (Nm) was the highest for the polar communities (Figure 2A). The iCAMP framework indicated that drift was the dominant factor shaping community assembly in the polar microbial communities (Figure 3A) [48]. The polar regions also have the highest proportion of neutral taxa. The seasonal freeze-thaw and extreme cold conditions are probably responsible for the lower diversity (Figure 1E, F), which in turn causes smaller community sizes [49]. As the overall community size decreases, the relative importance of drift increases (Figure 2A) [50].. The PERMDISP and the BCom analyses reveal a more homogeneous community structure caused by strong environmental filtering. Altogether, these analyses suggest that stochastic processes in polar regions operate within narrow environmental constraints. Environmental filtering limits which taxa can persist, while stochastic processes influence their relative abundances.

Stochastic processes played a more important role in temperate communities than polar communities, as evidenced by the R2 values, NCM analysis, and the NST values. The high BCom values and the dissimilarity of the temperate communities being higher than the null communities suggest high spatial turnover and environmental heterogeneity in temperate regions. The iCAMP results indicate notable roles of deterministic processes (homogeneous and heterogeneous selection) in shaping microbial community assembly in temperate regions, at the bin-level. Previous studies have also shown that homogeneous selection is a key contributor to community assembly [51]. This suggests that environmental selection and niche specialisation drive microbial community assembly. The absence of time series data limits the temporal resolution of our study, thereby reducing its ability to capture dynamic changes and temporal patterns within the microbial community.

Co-occurrence networks were constructed to investigate the ecological interactions and structural organisation underlying the observed patterns. The polar regions displayed the highest modularity of 0.778 across latitude zones (Table 1). Higher modularity is a characteristic of a more stable community, as perturbations affecting taxa in one module are less likely to affect the rest of the network [52]. The taxa acting as connectors and module hubs are specifically important for maintaining the network structure and stability. Polar microbial communities displayed the highest fraction of connectors and module hubs (Figure 6), suggesting that certain taxa in these communities play disproportionately important roles in maintaining the overall network stability. The natural connectivity analysis (Supplementary Figure 5) revealed that the polar communities are highly vulnerable to hub removal, as they exhibit the most negative slopes. While high modularity in the polar communities provides local stability, the high dependence on connector genera creates systemic vulnerabilities [53]. The taxa acting as connectors in the polar microbial co-occurrence network include Moritella [54] and Oleispira [55], which require low temperatures for survival and can degrade complex organic material. The increased vulnerability of polar communities due to increased modularity, along with the nature of the connector taxa, suggests that they may face risks from warming temperatures [56].

While the latitude-specific networks provided valuable insights into how community structure and stability vary across latitude gradients, it is also vital to understand the global ocean microbial co-occurrence network to investigate the interaction patterns and phylum-level organisation. This allowed for the improvement of the understanding of the ecological trends, such as dominant phyla, the nature of the inter- and intra-phyla interactions. Over 60% of the genera in the global ocean network belong to *Proteobacteria* and *Bacteroidota* (Figure 4A), consistent with findings from earlier ocean microbiome studies [17, 57–62]. *Proteobacteria* make up a large proportion of the nodes in the global ocean microbial networks due to their well-documented metabolic versatility and ecological adaptability [63, 64]. The high representation of *Bacteroidota* in the ocean microbial networks can be attributed to their ability to degrade complex carbohydrates, a trait enabled by their specialised polysaccharide utilisation loci (PULs) [65, 66]. The ocean microbiome had higher inter-phyla interactions compared to intra-phyla interactions (Figure 4B) across nodes. This aligns with studies which suggest that functional complementarity, rather than phylogenetic relatedness, governs ecological interactions in microbial communities [67, 68].

Specialists act as module hubs and connectors (Figure 7D), underscoring their role as keystone taxa in maintaining network structure and stability. Despite their widespread prevalence, the generalists act as peripherals, indicating that they interact broadly but weakly, or exist transiently in the community, as evidenced by the stochastic community assembly. To further understand the role of the specialists in network stability, the natural connectivity of the network was computed after removing a fraction of the specialists and generalists (Figure 7E). The specialists contribute more to the network stability than the generalists. The asymmetric vulnerability suggests that conservation of the specialist taxa is critical for sustaining marine microbial communities under perturbations such as warming and nutrient shifts and show that even rare taxa warrant conservation attention. Future research could utilise temporal data with species-level resolution to enhance understanding of marine microbiome resilience and functional dynamics under ongoing climate change.

## 5 Data Availability

The accession codes for the datasets used in the study are PRJEB42019, PRJEB36282, PRJEB36283, PRJEB25224, PRJNA656268, PRJNA1122929, PRJEB40762, PRJEB45011 and PRJEB27154.

## 6 Acknowledgements

The authors would like to thank Shradha Sharma and Pratyay Sengupta for their valuable insights and feedback on the manuscript. We are also grateful to Barathi L for her guidance and support in applying QIIME2 for data pre-processing. The authors used an AI-based language editing tool (ChatGPT 4.0 and Grammarly) to improve grammar and readability of the manuscript. No AI tools were used for data analysis, interpretation, or drawing scientific conclusions.

