## Supplementary Information for "Deciphering Global Patterns of Marine Microbial Community Assembly and Network Stability"

Indian Institute of Technology Madras

Chennai - 600 036, India

### Supplementary Information

### Supplementary Files

- **Supplementary File 1: Metadata of the samples**
- **Supplementary File 2: Denoising Parameters**
- **Supplementary File 3: Denoising Statistics**

### Supplementary Tables

**Supplementary Table 1.** Data used in this study. The number of samples taken from each study and the respective accession IDs are shown.

| **Project** | **Number of Samples** | **Relevant Project IDs** |
|---|---|---|
| Earth Microbiome Project | 1405 | QIITA IDs 10178, 678, 713, 723, 905, 1039, 1235, 1240 |
| Tara Oceans Project | 169 | PRJEB36282, PRJEB36283 |
| Malaspina Expedition | 320 | PRJEB25224, PRJEB45011, PRJEB27154 |
| Global Ocean Sampling | 47 | PRJEB40762 |
| Bio-GO-SHIP | 465 | PRJNA656268 |
| Australian Microbiome Project | 2369 | PRJNA1122929 |

**Supplementary Table 2: Multivariate Analysis of Microbial Beta Diversity Using PERMANOVA** The results show the $R^2$, which indicates the variation captured by each covariate, and the $p$-value indicates the significance of the results.

| **Factors** | $R^2$ **value** | $p$**-value** |
|---|---|---|
| Latitude Zones | 0.125 | 0.001 |
| Oceans | 0.119 | 0.001 |
| Sample Origin | 0.046 | 0.001 |
| Project | 0.081 | 0.001 |

**Supplementary Table 3: Canonical Correspondence Analysis of Marine Microbial Community Structure and Environmental Drivers** The table shows the correlation of environmental variables with the first two canonical axes (CCA1 and CCA2). It includes the coefficient of determination ($R^2$), indicating the strength of the relationship, and the statistical significance ($p$-value), indicating whether the observed correlation is significant.

| Variable | CCA1 | CCA2 | $R^2$ | $p$-value |
|---|---|---|---|---|
| Latitude Zone | -0.069 | -0.997 | 0.645 | 0.001 |
| Oceans | 0.255 | -0.966 | 0.090 | 0.001 |
| Sample Origin | 0.976 | -0.217 | 0.816 | 0.001 |

**Supplementary Table 4A: ANOVA-Based Comparison of Microbial Alpha Diversity Across Latitude Zones, Oceans, and Sample Origins** Results of ANOVA on the Shannon Diversity Index, showing the contribution of Latitude Zones, Oceans, and Sample Origin to observed diversity patterns across microbial communities.

| Factor | Df | Sum Sq | Mean Sq | F-value | Pr (>F) |
|---|---|---|---|---|---|
| Latitude Zones | 2 | 355.3 | 117.63 | 407.815 | $< 2 \times 10^{-16}$ |
| Oceans | 4 | 11.6 | 2.89 | 6.646 | $2.49 \times 10^{-5}$ |
| Sample Origin | 3 | 329.6 | 109.86 | 252.227 | $< 2 \times 10^{-6}$ |

**Supplementary Table 4B: ANOVA-Based Comparison of Microbial Alpha Diversity Across Latitude Zones, Oceans, and Sample Origins** The results of ANOVA on the Chao1 Richness Index indicate significant differences in species richness, as explained by Latitude Zones, Oceans, and Sample Origin.

| Factor | Df | Sum Sq | Mean Sq | F-value | Pr (>F) |
|---|---|---|---|---|---|
| Latitude Zones | 2 | 2820998 | 1410494 | 308.66 | $< 2 \times 10^{-16}$ |
| Oceans | 4 | 195740 | 48935 | 10.71 | $1.23 \times 10^{-8}$ |
| Sample Origin | 3 | 1799548 | 599849 | 131.26 | $< 2 \times 10^{-16}$ |

**Supplementary Table 5: Kruskal-Wallis test on the differentially abundant phyla in each latitude zone** The table shows phyla whose abundance distributions differ significantly between zones. The zone in which they are differentially abundant and the associated p-values are indicated.

| Phyla | $p$-value | Latitude Zone |
|---|---|---|
| *Campylobacterota* | $2.10 \times 10^{-230}$ | Temperate |
| *Bacteroidota* | $4.57 \times 10^{-190}$ | Polar |
| *Firmicutes* | $2.30 \times 10^{-158}$ | Tropical |

**Supplementary Table 6: Topological Roles of Genera in Latitude-Specific Co-occurrence Networks** Percentage of microbial genera classified as Peripherals, Connectors, and Module Hubs within the Polar, Temperate, and Tropical co-occurrence networks

| Group | Connectors | Module Hubs | Peripherals |
|---|---|---|---|
| Polar | 26.80% | 1.50% | 71.64% |
| Temperate | 2.22% | 0.98% | 96.82% |
| Tropical | 5.20% | 1.91% | 92.87% |

**Supplementary Table 7: Null model analysis on the PERMDISP results** This table illustrates the comparison between observed dispersion patterns and those expected under random assembly processes, helping to infer the ecological mechanisms driving community variability.

| Group | Dissimilarity (Observed) | Dissimilarity (Null) | F-value | ***p*-value** |
|---|---|---|---|---|
| Polar | 0.340 | 0.509 | 0.92 | $< 0.001$ |
| Temperate | 0.544 | 0.509 | 0.79 | $< 0.001$ |
| Tropical | 0.480 | 0.509 | 1.31 | $< 0.001$ |

### Supplementary Figures

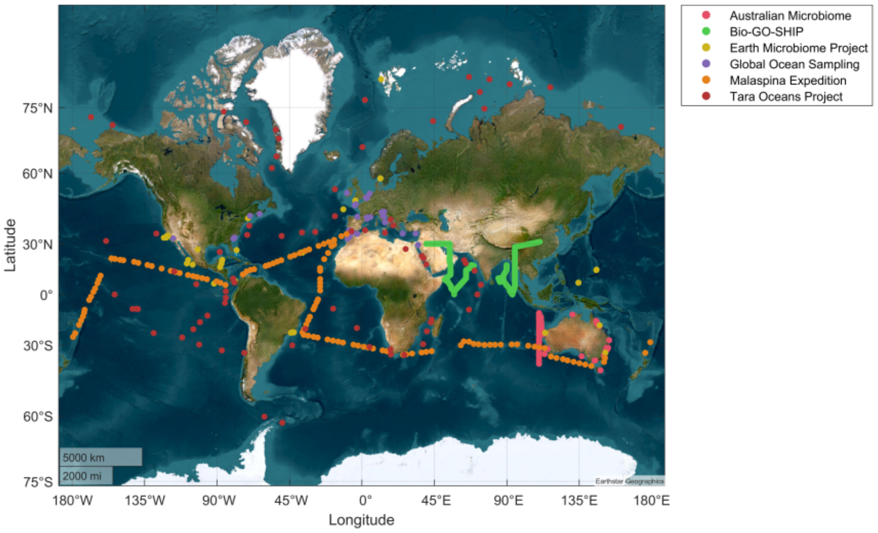


**Supplementary Figure 1: Geographic Distribution of the samples** The samples utilised in the study are geographically mapped and colour-coded based on the dataset from which it is obtained

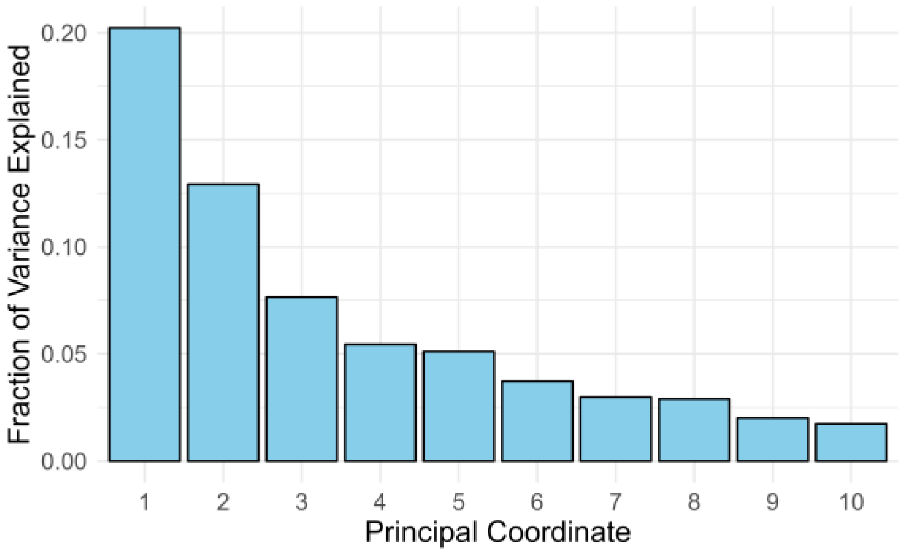


**Supplementary Figure 2: Scree plot depicting variation caused by each of the principal components .**

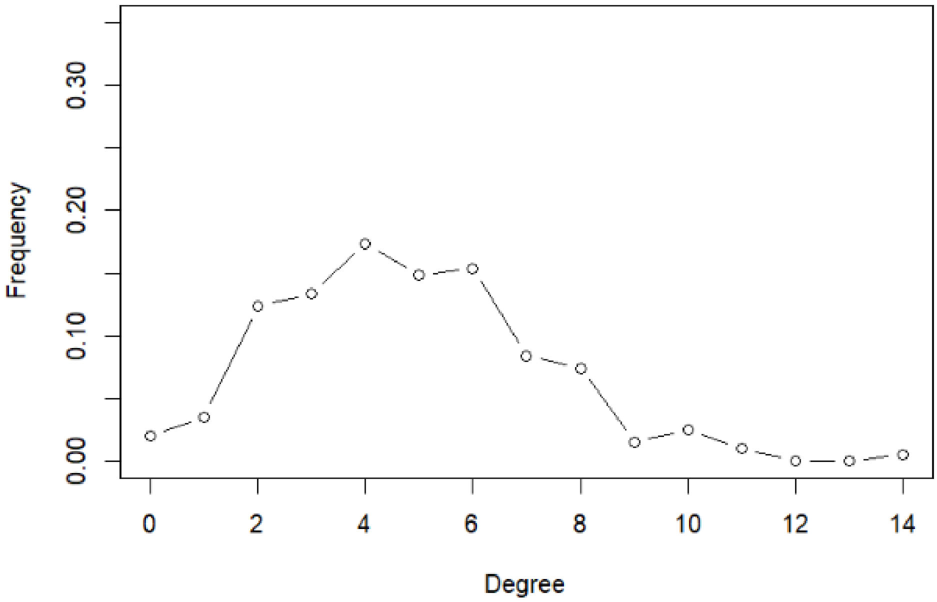


**Supplementary Figure 3A: The degree distribution of the latitude-specific co-occurrence networks** Degree Distribution of polar microbial co-occurrence networks.

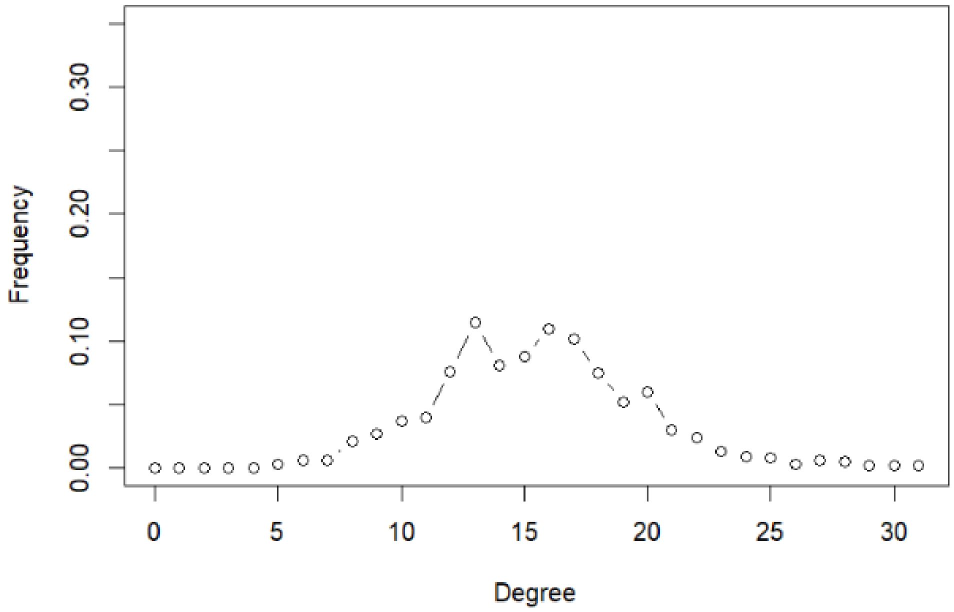


**Supplementary Figure 3B: The degree distribution of the latitude-specific co-occurrence networks** Degree Distribution of temperate microbial co-occurrence networks.

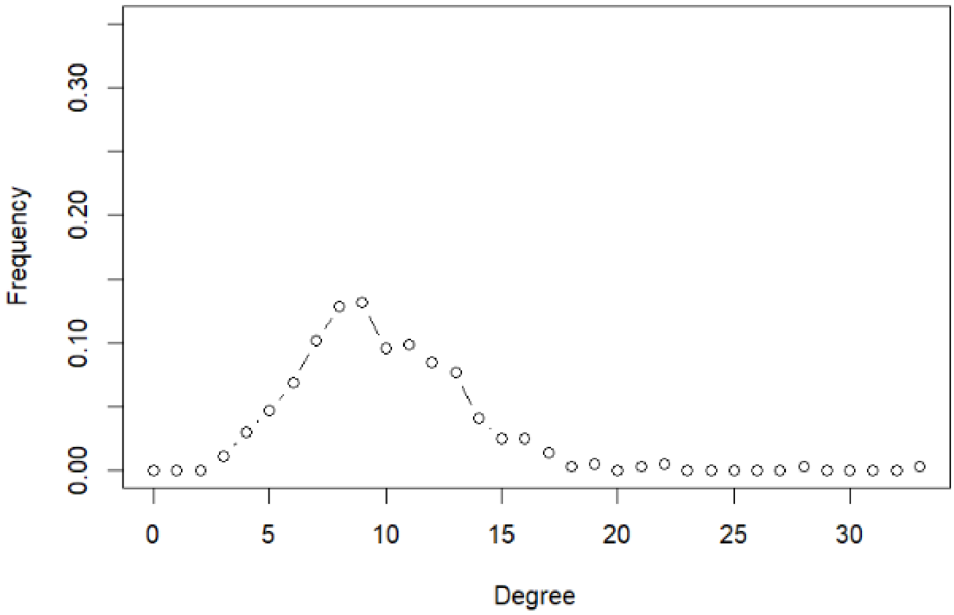


**Supplementary Figure 3C: The degree distribution of the latitude-specific co-occurrence networks** Degree Distribution of tropical microbial co-occurrence networks.

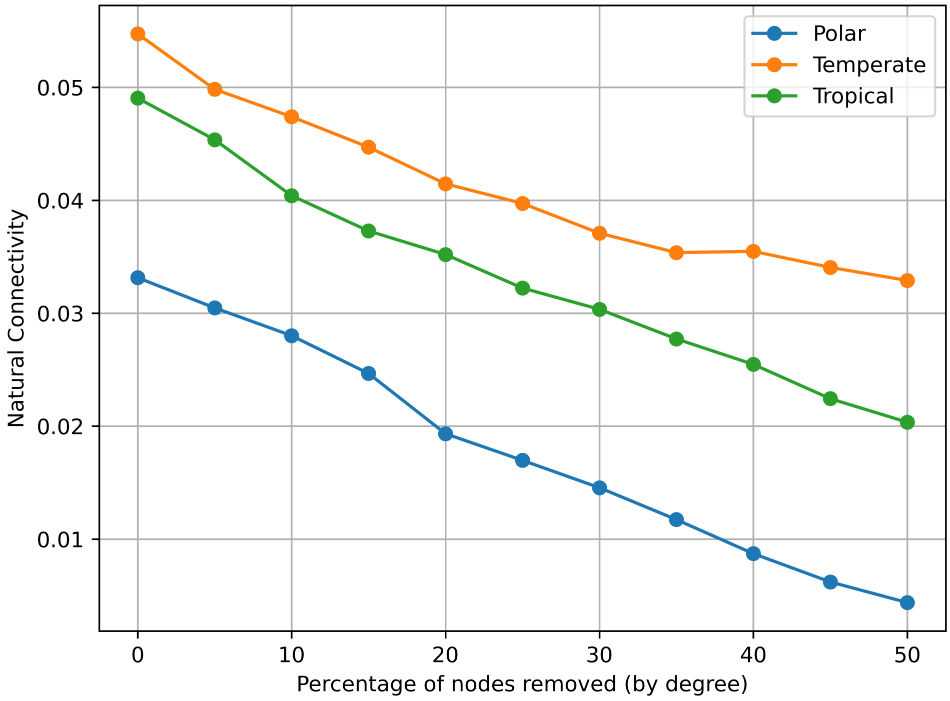


**Supplementary Figure 4: Natural Connectivity of the networks as nodes of the highest degree are removed sequentially** The variations in the natural connectivity after sequentially removing genera of the highest degree in polar, tropical and temperate networks (colour-coded).

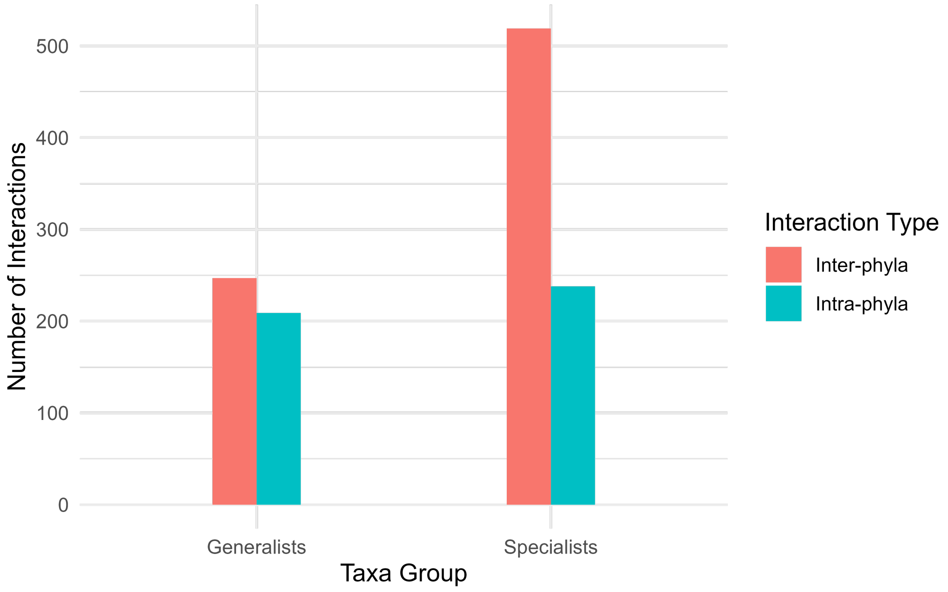


**Supplementary Figure 5: Number of inter- and intra-phyla interactions of the generalists and the specialists.**
